# Early Subclinical and Sex-Specific Cardiac Remodeling Precedes Heart Failure in Zebrafish with Human Actin T126I Mutation

**DOI:** 10.1101/2025.08.26.672352

**Authors:** Kendal Prill, John F. Dawson

## Abstract

Dilated cardiomyopathy (DCM) is a leading cause of heart failure with notable sex differences in disease susceptibility and progression. Mutations in sarcomeric proteins such as cardiac actin (ACTC1) contribute to familial DCM, yet the *in vivo* consequences of the ACTC1 p.T126I variant and its sex-specific effects remain unclear. We generated a zebrafish model carrying the orthologous Acta1b p.T126I mutation and conducted longitudinal analyses of cardiac phenotype, function, morphology, and gene expression stratified by sex. Mutants showed variable onset of cardiac dysfunction; surviving adults developed progressive DCM characterized by pericardial effusion, ventricular dilation, and reduced survival. Female mutants exhibited earlier and sustained diastolic dysfunction, more severe cardiac remodeling and significantly lower survival compared to males, revealing pronounced sexual dimorphism. Molecular profiling identified upregulation of the cardiac stress marker *nppb*, downregulation of hypertrophic transcription factors (*gata4, mef2ca*), stable or reduced sarcomeric gene expression, and sex-specific alterations in calcium handling genes (*serca2, pln1, slc8a1a*) and proteostasis regulators (*hsf1, bag3*). These findings demonstrate that the Acta1b p.T126I thin filament mutation drives progressive, sex-specific dilated cardiomyopathy in zebrafish, underscoring biological sex as a critical modifier of sarcomeric cardiomyopathy progression and providing a valuable platform for investigating sex-dependent mechanisms and developing targeted therapies relevant to women’s cardiovascular health.

## Introduction

Dilated cardiomyopathy (DCM) is a leading cause of heart failure and sudden cardiac death, characterized by ventricular dilation and impaired systolic function. Most DCM cases are idiopathic while others result from genetic mutations, environmental factors, or acquired injury [1–3]. Men tend to be more frequently and severely affected by DCM than women, suggesting sex-specific differences in disease susceptibility and progression [4]. Although over sixty genes have been linked to familial DCM, the underlying disease mechanisms remain incompletely resolved [5].

Primary mutations in sarcomere proteins, especially those affecting the thin filament, provide powerful entry points for dissecting core pathways of cardiac dysfunction. Among these, pathogenic variants in cardiac actin (ACTC1) though rare, are particularly informative because actin is essential for myofibril assembly, contractile force generation, and calcium regulation [6–9]. Additionally, ACTC1 mutations can produce either hypertrophic or dilated phenotypes, and affected individuals often exhibit variable disease onset and severity, with incomplete penetrance observed within the same family [10].

A c.383C>T mutation in human cardiac actin, ACTC1, is associated with familial dilated cardiomyopathy [10,11]. The cytosine-to-thymine transition leads to a threonine-to-isoleucine substitution at position 128 in the nascent actin protein, prior to cleavage of the two N-terminal amino acids, resulting in the p.T126I variant in the mature actin protein. *In vitro* assays determined the p.T126I variant causes reduced calcium sensitivity of thin filaments, reduced myosin ATPase activity and sliding velocity [12].

However, how this mutation triggers progressive heart failure *in vivo*, potential contribution of sex-specific disease pathogenesis, and the underlying molecular mechanisms are unclear.

Zebrafish are a powerful vertebrate model for cardiac research due to their genetic tractability, external fertilization, and short generation time, which enable detailed longitudinal studies of heart development and disease at embryonic, juvenile and adult stages [13,14]. Their cardiac structure, physiology, and gene expression profiles are highly conserved with humans, making zebrafish a valuable system to investigate human cardiomyopathies [15]. These features facilitate precise dissection of molecular and functional consequences of disease-causing variants such as the ACTC1 T126I variant in a living system.

Here, we generate and characterize zebrafish carrying the Acta1b (p.T126) mutation, orthologous to the human DCM-linked variant. By integrating longitudinal analyses of cardiac phenotype, function, morphology, and gene expression, we reveal that T126I induces variable, adult-onset DCM with striking sex-specific differences in remodeling and progression. These findings highlight how a thin filament mutation drives progressive dilated cardiomyopathy via sex-specific molecular remodeling, emphasizing sex as a key modifier of disease progression.

## Methods

### Generation of a Zebrafish Model of Human ACTC1-Dilated Cardiomyopathy

Zebrafish (*danio rerio*) have six cardiac actin paralogues that are expressed either constitutively or at specific developmental stages within the heart. There are three isoforms, *actc1a*, *actc1c*, and *acta1b*, predominantly expressed in cardiac tissue. *Actc1a* and *actc1c* are 100% identical at both the gene and protein sequence levels and share 99.47% sequence identity with human ACTC1 (Supplementary Fig. 1). In comparison, *acta1b* is a single-copy gene and exhibits 98.94% sequence identity with ACTC1 (Supplementary Fig. 1) [16]. Due to the redundancy of *actc1a/actc1c*, which may complicate functional analyses through genetic compensation and would require simultaneous targeting of both loci, *acta1b* represents a more tractable and practical model for introducing and studying human cardiac actin mutations in zebrafish [17]. The zebrafish line, *acta1b^hsc230^*, carrying the human c.383C>T (p.T126I) mutation in *acta1b* was generated in the AB strain background and validated by the Zebrafish Genetics and Disease Models Core, SickKids Research Institute (Supplementary Fig. 1).

### Zebrafish Husbandry, Line Maintenance and Ethics

The *acta1b^hsc230^* mutant strains were maintained as homozygotes and heterozygotes, along with the homozygous Tupfel Long Fin/AB strain (TLAB) wild-type strain, in the Hagen Aqualab at the University of Guelph. All zebrafish were housed in a recirculating aquatic system at 28.5°C, under a 12:12 hour light/dark cycle, in accordance with established zebrafish husbandry recommendations [18]. Embryos were obtained by crossing heterozygous adults; offspring were raised to adulthood for line maintenance and experimental analysis. All protocols were carried out following the guidelines stated by the Canadian Council for Animal Care and the University of Guelph (AUP#5144). Throughout this study, *acta1b^hsc230^* and *acta1b^hsc230^*^/+^mutants were compared to TLAB wild-type controls for every assessment and experiment. If an animal showed signs of stress, suspected pain or their quality of life had been significantly impacted by the progression of the disease, they were euthanized by submersion in an ice slurry for 45 mins until gill movement ceased and they did not respond to touch.

### Genotyping *acta1b^hsc230^* mutants

*Acta1b* exons were amplified from genomic DNA extracted from homozygous and heterozygous *acta1b^hsc230^* larvae and adult fin clips using *acta1b* FWD: GGGTATCCTCACTCTGAAATA and REV: CGGTTGTGACGAAAGAATAA primers. PCR products were purified and submitted for Sanger sequencing through the University of Guelph Agriculture and Food Laboratory Service. Sequencing chromatograms were analyzed using 4Peaks software to distinguish wild-type, heterozygous, and homozygous *acta1b^hsc230^*genotypes.

### Frailty Assessment of Adult Zebrafish

Fish were netted and allowed to acclimatize in a 1.1 L tank with a divider to ensure primarily lateral movement. Tests were recorded using an iPhone Xs camera (Apple Inc, Cupertino, USA) positioned 1.5 feet from the tank on the same table. Two min of swimming was used to assess the speed (20 inches in 15 seconds) and casual endurance of individual fish. This was followed by a stimulation, and 1 min recovery repeated 3 times total. Lastly, fish food pellets (Gemma 300; Skretting, Canada) were added to the front of the tank when the fish was at the back. Response to food was timed based on 3 tiers: 15 seconds, 30 seconds and 1 min. Additional test categories were normal anatomical appearance, swimming horizontal and level, and continuous swimming. All assessments were based on a binary system where positive scored 1 point and absence or negative responses scored 0. 6-month-old wild-type and *acta1b^hsc230^* fish were excluded from the frailty assessment due to signs of stress not observed in the 24-27-month-old cohort.

### Echocardiography and Analysis

Adult zebrafish were anesthetized using a combination of MS-222 (tricaine; TCI America, USA) and isoflurane (TCI America, USA) [19]. Cardiac function in adult zebrafish was acquired and assessed at 4, 6, and 24-27 months of age using a high-frequency ultrasound imaging system (Vevo 3100 LT, FUJIFILM Visual Sonics; Canada) as previously described [20]. Ultrasound images and measurement of cardiac parameters were performed using Vevo 3100 imaging software (VEVO Lab, FUJIFILM Visual Sonics Inc., 2021) as previously described [20]. Averages of wild-type and heterozygous individuals were done due to limited tank space to house all fish individually during assessments. For 24-27-month-old fish: wild-type males n=11, wild-type females n=11, *acta1b^hsc230/+^* males n=8, *acta1b^hsc230/+^* females n=6, *acta1b^hsc230^*males n=13, *acta1b^hsc230^* females n=9. For 6-month-old fish: wild-type males n=12, wild-type females n=14, *acta1b^hsc230^*males n=13, *acta1b^hsc230^* females n=14.

### Heart Dissection and Physiological Assessment

Adult zebrafish were euthanized by submersion in an ice slurry for 45 mins until cessation of gill movement and lack of response to stimuli. Six-month-old fish were freshly dissected after euthanasia while 19-27-month-old fish were fixed in formalin (Fisher Scientific, Canada) at 4°C. Before dissection of the thoracic cavity, the length (tip of mouth to caudal peduncle) and weights of all adult zebrafish were recorded. Lateral and ventral images were taken of individuals using the iPhone Xs camera (Apple Inc; Cupertino, USA) freehand and mounted to a Zeiss SteREO Discovery V8 microscope (Oberkochen, Germany). A sagittal cut followed by two transverse cuts, to form a window, were made on the ventral surface of the thoracic region to allow visualization of the heart before removal [21]. Distal regions of the bulbus arteriosus and sinus venosus were cut to release the heart. The hearts were imaged, measured and weighed before proceeding to downstream experimental analysis. For 16-27-month-old fish: wild-type males n=21, wild-type females n=15, *acta1b^hsc230/+^* males n=8, *acta1b^hsc230/+^* females n=4, *acta1b^hsc230^*males n=29, *acta1b^hsc230^* females n=17. For 6- month-old fish: wild-type males n=10, wild-type females n=14, *acta1b^hsc230^* males n=11, *acta1b^hsc230^* females n=13.

### Cryosectioning, Histology and Immunofluorescence

Hearts designated for histological analysis were washed 3 x 10 min phosphate buffered saline (PBS), washed in a series of sucrose baths, and washed again in tissue freezing medium (Epredia, USA) [22]. Hearts were sectioned at 12 um thickness on a cryostat (Leica CM3050 S; Canada) and baked onto Superfrost Plus slides (Fisher Scientific, Canada) for 2 hours at room temperature.

Heart sections were sent to the University of Guelph Animal Health Laboratory Histology Suite for Hematoxylin and Eosin staining. Stained heart sections were imaged using a mounted iPhone X camera (Apple Inc; Cupertino, USA) mounted to a Carl Zeiss Jena compound microscope (Zeiss).

For antibody staining of sectioned tissue, cryosections were hydrated for 3 mins in PBTD (PBS, 0.1% tween-20, 1% DMSO) in square petri plates. Antibody staining was performed as previously described with 1/500 dilution of Rhodamine Phalloidin (Cytoskeleton, USA). Stained tissue was imaged using Diskovery Spinning Disk confocal (Quorum, Canada) and images processed using Volocity software (Quorum, Canada). For all staining analysis: male wild-type and *acta1b^hsc230^* n=2 each; female wild-type and *acta1bhsc230* n=3 each).

### RNA Extraction and Quantitative PCR

Freshly dissected 6-month-old hearts were flash frozen in liquid nitrogen before RNA extraction. RNA was extracted from flash frozen hearts using TRIzol-chloroform method as previously described [23]. mRNA was normalized to 100 ng/ul prior to cDNA synthesis (5X All-In-OneBioBasic RT PCR Mastermix; BioBasic, Canada). All qPCR was performed using GB-Amp InFluor Green qPCR Mix (GeneBio Systems Inc, Canada), primers (Supplementary Table 1) and Applied Biosystems QuantStudio 7 Pro Real-time PCR System (Thermo Fisher Scientific, USA). Relative fold change in expression for each gene was determined using the ΔΔCt method [24]. Ct values were first normalized to the reference gene (*rpl13a*), then compared to the same-sex wild-type control group, which was set to a value of 1. Both male and female wild-type n=3 each; male *acta1bhsc230* n=4; female *acta1b^hsc230^* n=5).

## Results

### Distinct Phenotype Manifestations, Variable-Onset Mortality and Sex-Specific Survival are Observed in Zebrafish Carrying the Human T126I Cardiac Actin Mutation

In humans, a c.383C>T (p.T126I) mutation in cardiac actin (ACTC1) has been associated with dilated cardiomyopathy [10]. Our prior *in vitro* work demonstrated that the ACTC1 T126I variant reduces thin filament calcium sensitivity, myosin ATPase activity, and actomyosin sliding velocity [12]. To explore *in vivo* consequences, we investigated phenotypic and molecular changes in zebrafish harboring the orthologous T126I mutation (*acta1b^hsc230^*).

Zebrafish carrying one or two copies of *acta1b^hsc230^* exhibited a spectrum of cardiac phenotypes with variable onset. A subset of mutant larvae displayed severe early cardiac dysfunction characterized by bradycardia, pericardial and body edema, heart degeneration, and absence of swim bladder inflation, resulting in lethality between 4 and 7 days post-fertilization (Supplementary Fig. 2A-C). A large proportion of mutants survived larval stages without overt abnormalities and are the focus of this study to characterize the adult-onset cardiac phenotypes and disease pathway.

Adult homozygous and heterozygous mutants appeared morphologically normal until ∼6 months of age, after which the incidence of phenotypic abnormalities and mortality increased notably around 19 months (Fig. 1, Supplementary Fig. 2D-H). Body lengths and mass indices were comparable between most groups, except homozygous females who were significantly heavier than wild-type females and homozygous males (Supplementary Fig. 3A&B). Phenotypes were variable among and within sex and genotype groups, consistent with incomplete penetrance and variable expressivity seen in cardiac actinopathies in humans [5,25,26].

**Figure 1.**
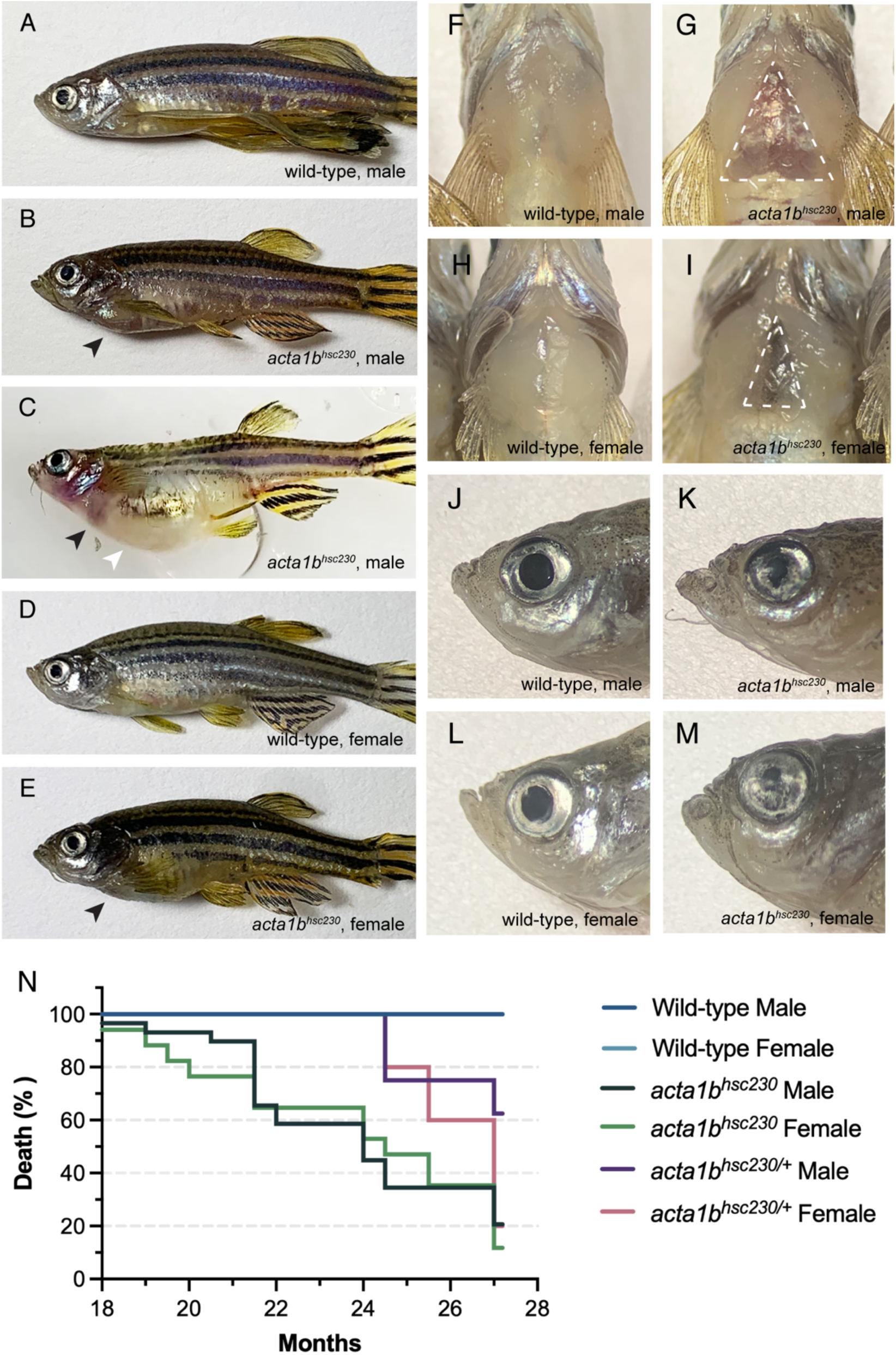
Adult-onset morphological abnormalities and reduced survival in zebrafish carrying the *acta1b^hsc230^* mutation. (A-E) Lateral views of adult zebrafish at 24-27 months of age. Wild-type male and female zebrafish exhibit normal morphology (A, D). Homozygous *acta1b^hsc230^* mutant males and females display progressive phenotypic severity, including pericardial effusion (B,C&E; black arrowheads) and occasional generalized edema (C; white arrowhead). (F-I) Ventral views of wild-type and *acta1b^hsc230^*mutants further highlight pericardial effusion and ventral cardiac protrusion in mutant males and females (dotted white triangle). (J-M) A subset of adult mutants exhibit reduced pupil size, predominantly in the left eye. (N) Longitudinal analysis of survival in *acta1b^hsc320^* mutant zebrafish.

Both heterozygous and homozygous mutants of both sexes developed pericardial effusion correlating with ventral protrusion of the heart (Fig. 1B,C,E-I; Supplementary Fig. 3C) [27]. Some homozygous males exhibited severe generalized body edema mimicking gravid females (Fig. 1C; Supplementary Fig. 2H). Remarkably, some adult mutants displayed progressive pupil size reduction, predominantly in the left eye, days to weeks before cardiac symptoms (Fig. 1J–M, Supplementary Fig. 3C). Additional abnormalities such as spontaneous hemorrhaging, gasping-like behavior and difficulty swimming were also observed (Supplementary Fig. 3C). Using an adapted frailty assessment analogous to human cardiac patients [28], mutants scored significantly lower and showed reduced feeding responsiveness, impaired swimming, and increased stationary periods (Supplementary Fig. 3D).

Survival tracking until 24.5 or 27 months revealed sex- and genotype-dependent differences (Fig. 1N): survival was lowest in homozygous females (11.7%) and males (20.7%), while heterozygous males showed relatively higher survival (62.5%). These results demonstrate that zebrafish with the T126I cardiac actin mutation exhibit heterogeneity in disease onset, severity, and survival, with female mutants displaying the poorest outcomes. The nature and progression of the mutant phenotype suggests underlying cardiac functional deficits.

### Cardiac Dysfunction in Aged and Young Adult *acta1b^hsc230^* Mutants

Given the variation in mutant phenotypes, and the greater mortality observed in female mutants, we wanted to determine if cardiac function is compromised in the cardiac actin mutants. High-frequency echocardiography of 24.5- and 27-month-old adults revealed pronounced pericardial edema and altered cardiac morphology in mutants compared to wild-type controls (Fig. 2A–C). Most mutant groups, except heterozygous males, exhibited significantly lower heart rates, and reduced ejection fraction, consistent with impaired systolic function (Fig. 2D&E) [27]. Cardiac output was markedly decreased in homozygous females (Fig. 2F), likely due to their increased body surface area (Supplementary Fig. 3B).

**Figure 2.**
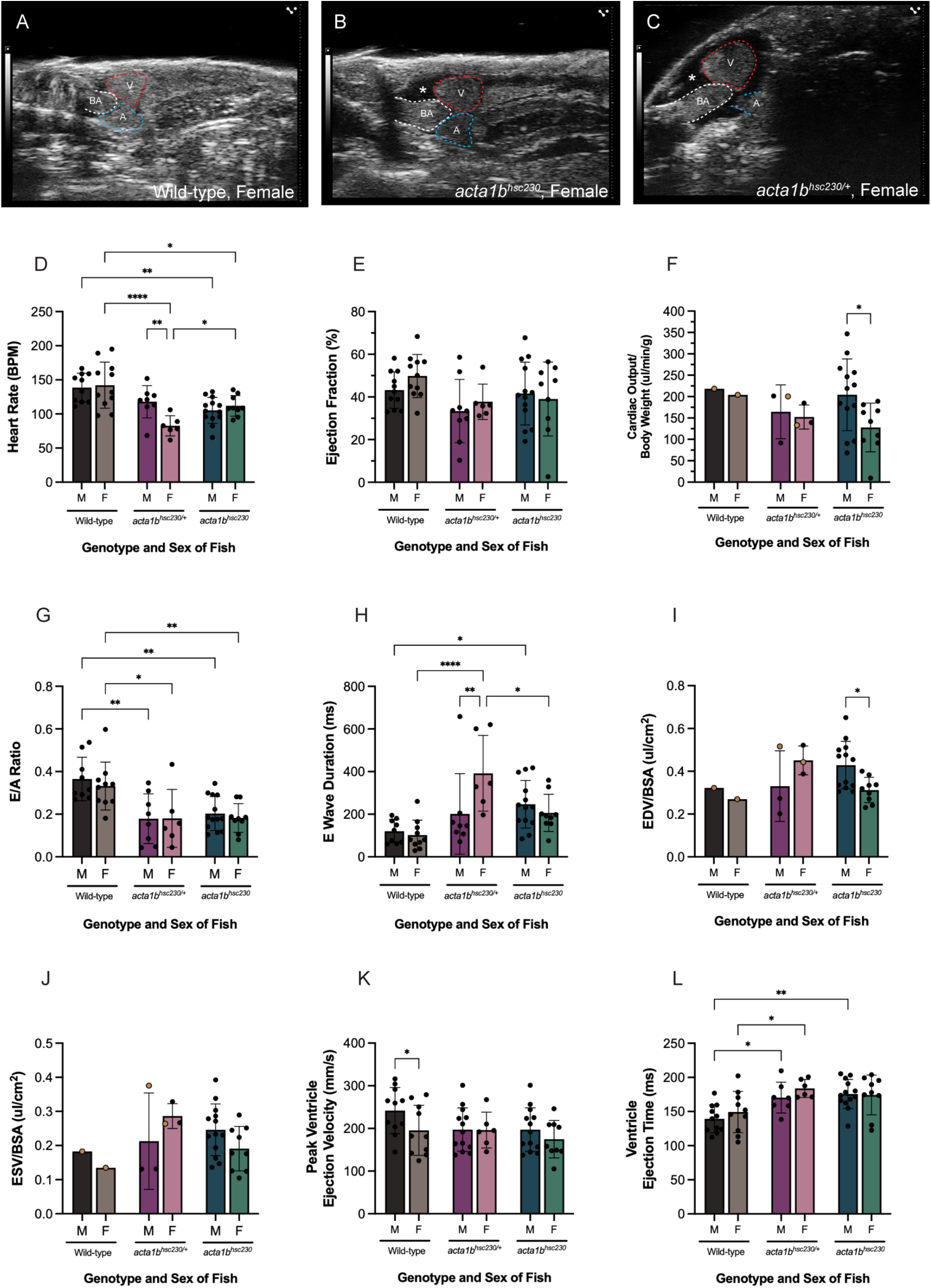
Aged *acta1b^hsc230^* mutants exhibit diastolic and systolic dysfunction. (A-C) Representative high-frequency ultrasound images of hearts from wild-type and *acta1b^hsc230^* mutant zebrafish. Mutant fish display pericardial edema (asterisk). Ventricular (V, red dotted line), atrial (A, blue dotted line), and bulbus arteriosus (BA, white dotted line) boundaries are indicated in panels A–C. Quantitative comparison of cardiac functional parameters among wild-type, heterozygous (*acta1b^hsc230^*^/+^) and homozygous (*acta1b^hsc230^*) mutants at late adult stages: (D) heart rate (beats per minute, BPM), (E) ejection fraction, (F) cardiac output normalized to body weight, (G) E/A ratio, (H) E wave duration, (I) end diastolic volume (EDV) normalized to body surface area (BSA), (J) end systolic volume (ESV) normalized to BSA, (K) peak ventricular ejection velocity, and (L) ventricular ejection time. Individual data points are shown as black dots; group means are shown as yellow dots. Data are presented as mean ± SD. *, p< 0.05 by two-way ANOVA with Tukey’s post hoc multiple comparisons.

Both heterozygous and homozygous *acta1b^hsc230^* mutants exhibited significantly reduced ratios of early (E) and atrial (A) peak ventricular inflow velocities (E/A ratio) and prolonged early ventricular filling phases, indicative of diastolic dysfunction (Fig. 2G&H) [19]. End diastolic and end systolic ventricular volumes were elevated across all mutant groups, suggesting impaired contractile clearance and increased preload (Fig. 2I&J) [27]. Despite the ability of mutant ventricles to reach wild-type ejection velocities, contraction times were significantly prolonged, especially in males (Fig. 2K&L). Collectively, these parameters indicate combined systolic and diastolic impairment reminiscent of dilated cardiomyopathy.

To determine whether the pronounced cardiac abnormalities in older adult T126I mutants were preceded by subtler cardiac changes, we performed echocardiography on homozygous *acta1b^hsc230^* zebrafish at 4 and 6 months. Pre-symptomatic, sex-specific abnormalities were apparent at 4 months, with reduced E/A ratio in females, and prolonged filling time and lower heart rates in both sexes (Fig. 3A-C). By 6 months, most measures were comparable to wild-type, except for persistently low E/A ratios in females (Fig. 3D-F). These findings demonstrate that functional impairment arises months before anatomical remodeling or heart failure becomes phenotypically evident.

**Figure 3.**
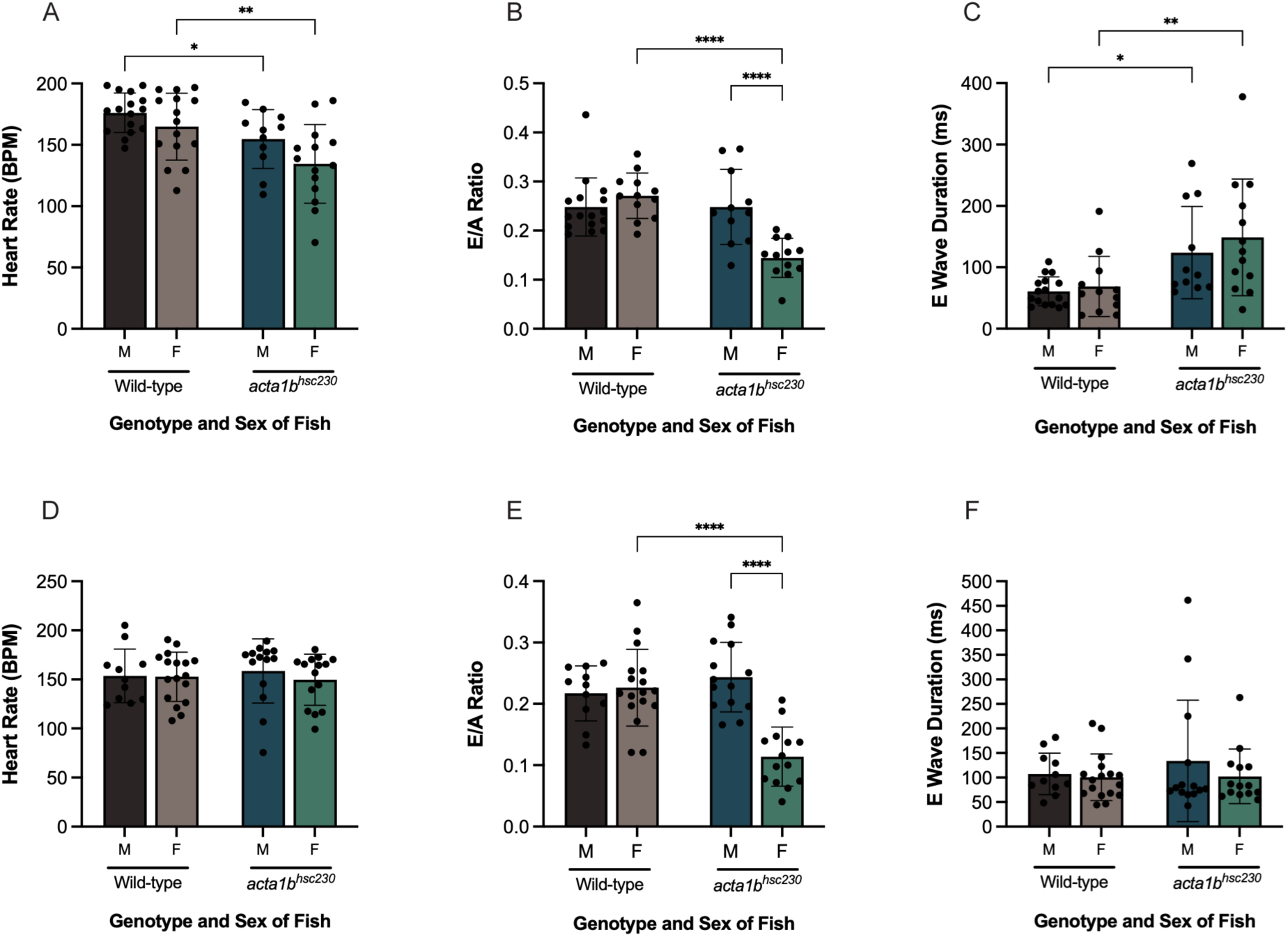
Young Adult *acta1b^hsc230^* fish present with diastolic dysfunction in sex-specific manners. Echocardiographic assessment of cardiac function in wild-type and *acta1b^hsc230^* mutants at 4 months of age, showing (A) heart rate, (B) early-to-atrial (E/A) peak ventricular inflow velocity ratio, and (C) E wave duration. Serial measurements of these parameters at 6 months of age are shown in (D) heart rate, (E) E/A ratio, and (F) E wave duration. Black dots represent individual fish. Data are presented as mean ± SD. *, p< 0.05 by two-way ANOVA with Tukey’s post hoc multiple comparisons.

### Late-onset Cardiac Remodeling and Myofibrillar Disorganization in ***acta1b^hsc230^*** Mutants

To assess whether the functional impairments were associated with morphological cardiac changes, we examined gross heart anatomy at early and late adult stages. At 6 months, mutant and wild-type hearts appeared normal (Fig. 4A&B). However, hearts from mutants aged 24.5 and 27 months showed variable remodeling, including atrial enlargement, ventricular distortion, bulbus arteriosus enlargement, and callus-like structures on the ventral face of the ventricle (Fig. 4C-F). Heart weight and size were elevated in homozygous mutants and to a lesser extent in heterozygotes (Fig. 4G&H).

**Figure 4.**
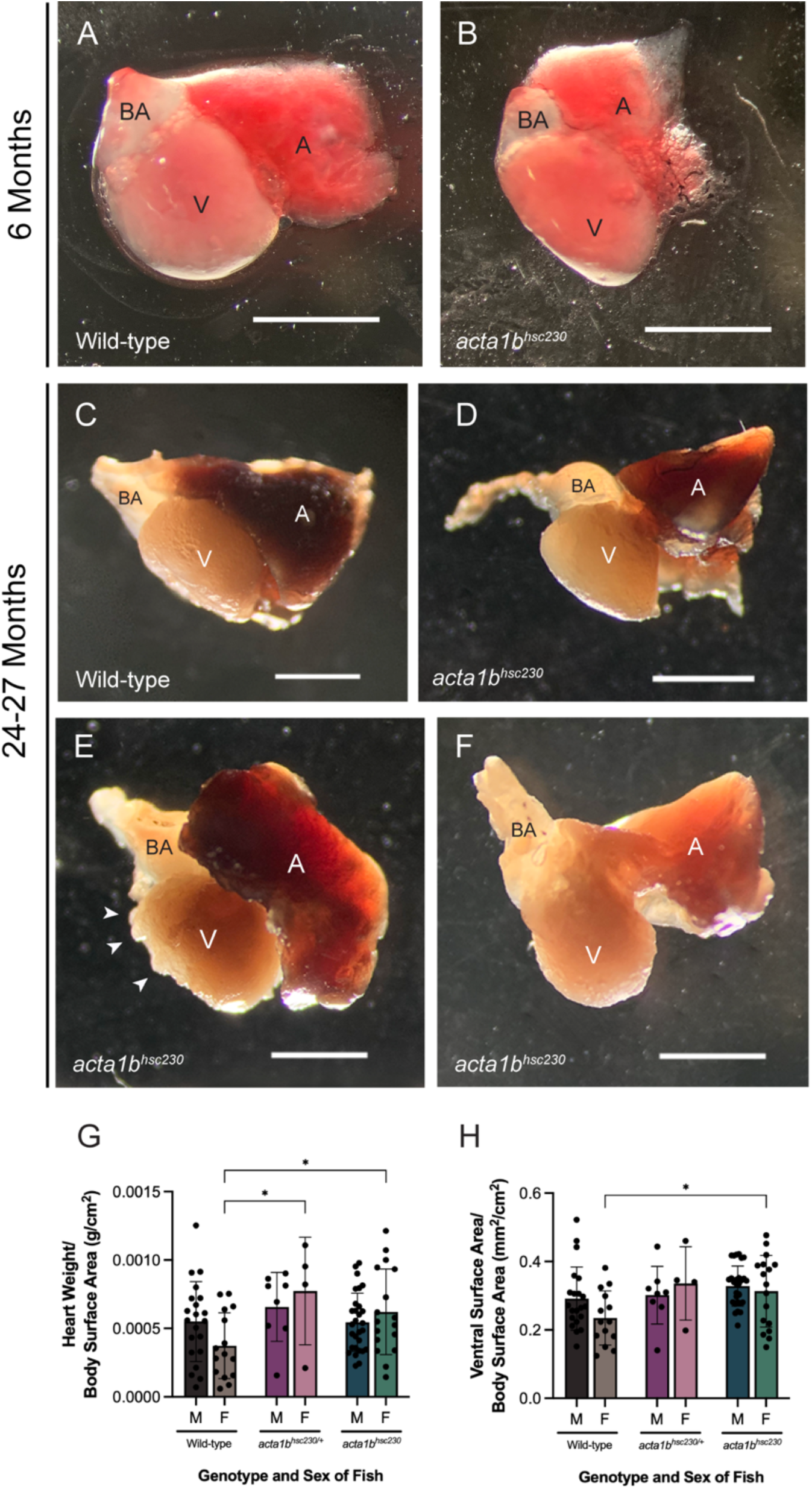
Comparison of wild-type and *acta1b^hsc230^* heart morphology at young and late adult stages. Freshly dissected hearts from wild-type (A) and *acta1b^hsc230^*mutant (B) adult zebrafish at 6 months of age. Fixed hearts from wild-type (C) and *acta1b^hsc230^* mutants at 24–27 months of age showing morphological differences in the bulbus arteriosus (BA), ventricle (V), and atrium (A). Scale bars represent 1 mm. Quantitative analysis of (G) heart weight normalized to body surface area and (H) area of the ventral ventricular surface normalized to body surface area for 24–27 month-old fish. Black dots represent individual animals; data are shown as mean ± SD. *, p< 0.05 by two-way ANOVA with Tukey’s post hoc test.

Histological analysis of late-stage mutant hearts revealed regions of trabecular disorganization and reduced cardiomyocyte thickness (Fig. 5A-F, asterisks). Actin immunofluorescence confirmed pronounced myofibrillar disarray in affected regions compared to medial and wild-type trabeculae (Fig. 5G-I). The absence of these structural defects in 6-month hearts (data not shown) suggests that progressive myofibrillar loss underlies the severe late-onset cardiac dysfunction.

**Figure 5.**
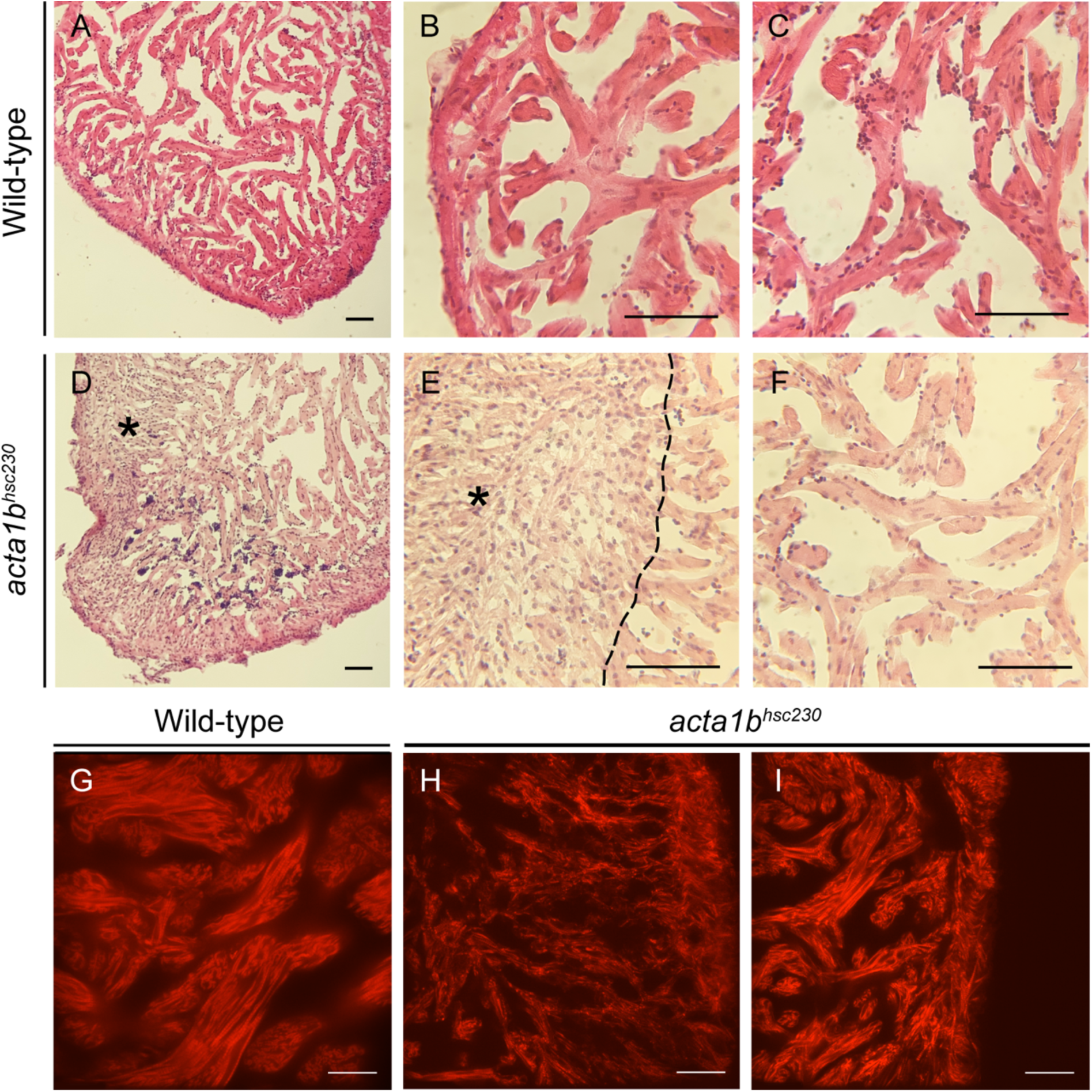
Myofibrillar loss and disorganization in *acta1b^hsc230^* mutant hearts at late adult stages. Hematoxylin and eosin staining from wild-type (A-C) and *acta1b^hsc230^* (D-F) hearts. The asterisk denotes a region of myofibrillar loss and the dashed line demarcates this region from normal trabeculae. Representative actin immunofluorescent images showing organization in wild-type trabeculae (G), disorganized regions (H) and normal-appearing trabeculae (I) in *acta1b^hsc230^* hearts. Scale bars: 50 um (A-F), and 33 um (G-I).

### Sex-specific Transcriptional Alterations in ***acta1b^hsc230^*** Mutant Hearts

To evaluate whether cardiac phenotypes observed in *acta1b^hsc230^*zebrafish mutants correspond to dilated cardiomyopathy (DCM) at the molecular level, we measured expression of established DCM markers. The natriuretic peptide gene *nppb*, a canonical marker of cardiac stress and remodeling [29,30], was significantly upregulated in mutants (Fig. 6A). Conversely, key cardiac transcription factors *gata4* and *mef2ca*, which mediate hypertrophic responses [31–33], were downregulated, consistent with impaired adaptive signaling typical of DCM. Sarcomere genes *amhc*, *vmhc, vmhcl, tnni1b, tnnt2a* and *tpm4a* showed stable or reduced expression (Fig. 6B)(mutant males had one individual outlier with significantly increased *vmhc* expression (Supplementary Fig. 4A)), contrasting with their upregulation during hypertrophic growth, further supporting a dilated remodeling profile.

**Figure 6.**
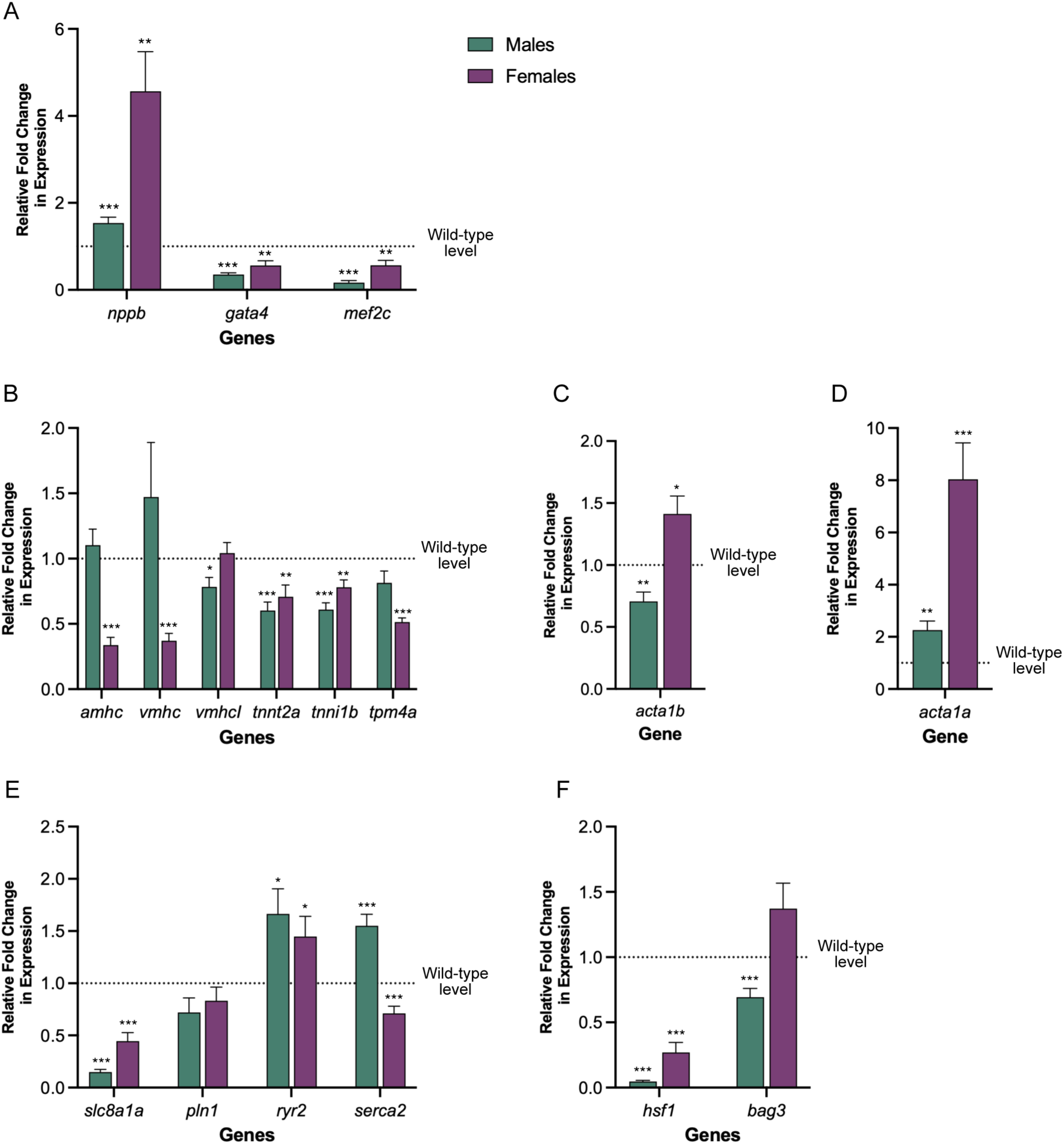
Sex-specific transcriptional alterations in *acta1b^hsc230^* mutant hearts at early asymptomatic onset of diastolic dysfunction. Relative mRNA expression levels of cardiac stress markers and remodeling-associated genes in male and female *acta1b^hsc230^* mutants at 6 months of age. Expression of (A) *nppb, gata4*, and *mef2ca*; (B) sarcomere genes *amhc, vmhc, vmhcl, tnnt2a, tnni1b*, and *tpm4a*; (C) *acta1b*; (D) *acta1a*; (E) calcium handling genes *slc8a1a, pln1, ryr2*, and *serca2*; and (F) proteostasis regulators *hsf1* and *bag3*. Expression values were normalized to sex-matched wild-type controls, indicated by the dotted line in each graph. Data are presented as mean + SEM. *, p<0.05 by t-test with unequal variance comparing mutants to wild-type within each sex.

Given the potential for genetic compensation among zebrafish cardiac actin paralogues [34], we assessed the expression of all six paralogues. We found that female mutants showed increased expression of *acta1b*, while males exhibited a decrease (Fig. 6C). *Acta1a* was significantly upregulated in all mutants, suggesting a possible compensatory response (Fig. 6D). Other actin paralogues, *actc1a/c and actc2,* showed lower or stable expression while *actc1b* showed variable changes between sexes (Supplementary Fig. 4B).

Since previous *in vitro* data indicated that the actin T126I mutation reduces calcium sensitivity [12], we investigated the expression of genes in the calcium handling pathway. Both sexes exhibited downregulation of *slc8a1a* (NCX1, sodium-calcium exchanger) and *pln1* (phospholamban), which regulates SERCA activity (Fig. 6E) [35,36]. Ryanodine receptor 2 (*ryr2b*) expression was elevated for both sexes, potentially increasing the release of more calcium from the sarcoplasmic reticulum [37]. However, *serca2* expression was differentially regulated between sexes, decreasing in females but increasing in males (Fig. 6E), suggesting sex-specific compensatory mechanisms in calcium homeostasis in response to desensitized thin filaments.

Considering that Acta1b p.T126I affects the function of thin and thick filaments of the sarcomere, we wanted to determine if any genes from the stress pathway were dysregulated. *Hsf1*, a master regulator of heat shock proteins [38,39], was downregulated in both sexes (Fig. 6F). However, the expression of co-chaperone *bag3*, essential for sarcomere maintenance and autophagy [40,41], was slightly elevated in females but decreased in males (Fig. 6F), highlighting sex-dependent differences in proteostasis and sarcomere quality control.

## Discussion

We have established a zebrafish model carrying the orthologous T126I cardiac actin mutation linked to familial dilated cardiomyopathy (DCM) in humans, providing new *in vivo* insights into sarcomeric DCM pathogenesis and progression. Through comprehensive longitudinal characterization of both male and female zebrafish, we demonstrate that this thin filament mutation induces progressive, variable-onset cardiac dysfunction with notable sexual dimorphism in disease severity, survival, and remodeling.

Marked sexual dimorphism emerged, with female mutants exhibiting more severe disease progression, reduced survival, and sustained early diastolic dysfunction compared to males, who showed partial functional recovery at younger adult stages (Fig. 1-3, Supplementary Fig. 2&3). This contrasts somewhat with epidemiological data indicating male predominance in DCM but aligns with emerging evidence that sex influences molecular remodeling and disease trajectory differentially [3,42]. Our findings highlight the importance of investigating sex-specific pathogenesis to better understand cardiomyopathy progression and therapeutic response.

Morphological analyses showed that *acta1b^hsc230^* mutant hearts are grossly normal in early adulthood but develop late-onset remodeling characterized by atrial enlargement, ventricular distortion, and trabecular disorganization (Fig. 4). Histological and immunofluorescence data indicate focal cardiomyocyte atrophy marked by myofibrillar loss rather than overt cell death at older stages (Fig. 5), consistent with compensatory dedifferentiation or regenerative attempts. Given the zebrafish’s notable regenerative capacity [42], this remodeling likely reflects maladaptive or insufficient repair contributing to cardiac dysfunction and heart failure.

Transcriptional profiling supports these phenotypes. Upregulation of *nppb* confirms cardiac stress and remodeling (Fig. 6) [29,30]. The downregulation of hypertrophy-associated transcription factors (*gata4* and *mef2ca*) and stable or reduced expression of sarcomeric genes distinguish the observed molecular phenotype from hypertrophic cardiomyopathy, consistent with DCM remodeling. The differential regulation of cardiac actin paralogue expression (Fig. 6; Supplementary Fig. 4) suggests partial genetic compensation with potential sex-dependent modulation that influences the severity of compensation. It is not clear why female mutant hearts would increase the expression of *acta1b^hsc230^* in contrast to male mutant hearts that decrease the expression of the mutant paralogue.

Notably, altered expression of calcium handling genes, especially sex-specific changes in *serca2,* upregulation of *ryr2* and downregulation of *pln1* and *slc8a1a*, suggest disrupted calcium homeostasis contributes to the phenotype (Fig. 6). This manipulation of calcium pathway genes suggests attempts to increase cytosolic calcium concentration, aligning with our prior demonstration that Actin p.T126I reduces thin filament calcium sensitivity. Resulting calcium overload in cardiomyocytes may impair ventricular relaxation and thus underlie the diastolic dysfunction seen in mutant animals. These disturbances are likely to trigger adaptive and maladaptive signaling cascades.

Furthermore, dysregulation of proteostasis pathways, including reduced *hsf1* and sex-dependent changes in *bag3* (Fig. 6), implicate impaired sarcomere maintenance and autophagic quality control in progressive dysfunction.

Mechanistically, the T126I variant impairs sarcomeric contractility primarily via reduced thin filament calcium sensitivity, leading to compromised force generation. This insufficiency may elicit compensatory increases in cytosolic calcium to maintain contraction, but chronic calcium overload can disrupt cardiomyocyte electrical and mechanical properties, induce mitochondrial damage, and activate maladaptive signaling [43–45]. These processes promote progressive myocardial remodeling, sarcomere disarray, and ultimately heart failure, as observed in our *acta1b^hsc230^* mutants.

Taken together, our findings identify critical molecular and functional pathways driving sarcomeric DCM with sex-specific modulation with the *acta1b^hsc230^ mutation*. This zebrafish model provides a valuable platform for dissecting early and late disease stages, elucidating sex as an important modifier, and testing targeted interventions to mitigate progressive heart failure in sarcomeric cardiomyopathies.

