## Supplementary Data for "Early Subclinical and Sex-Specific Cardiac Remodeling Precedes Heart Failure in Zebrafish with Human Actin T126I Mutation"

Supplementary Table 1. qPCR Primers Used to Quantify Relative Expression in *acta1b*$^{h230}$ Mutants.

| Gene | Forward Primer (5' → 3') | Reverse Primer (5'→ 3') |
|---|---|---|
| *rpl13a* | CTCTCAAGATTGTGCGTCTG | GGTGATGGCCTGGTACTT |
| *bnp/nppb* | CATGGGTGTTTTAAAGTTTCTCC | CTCAA TATTTGCCGCCTTTAC |
| *gata4* | TGTCAGACTACCACAACAACTC | GTGTCTGAATGCCCTCTTTCT |
| *mef2ca* | GAGAAGTGATGGGCGGATAC | CATGGAGCCCAGATGAAGAG |
| *amhc* | TCTGGAGCAAACCGAAAGAG | GATCCGACTCTTGCTTCTTCTT |
| *vmhc* | AAGAGCTGACATTGCAGAG | TCCACTTGAGCTTTACTCTTG |
| *vmhcl* | TGCTGAATCCCAAGTGAAC | TTCAGAAGATTTCCAGGACTTT |
| *tnnt2a* | CTGAGGAACAGAGTCAG | TGTGATTTGTGAGGATTC |
| *tnni1b* | GTCTGGAATGGAAGGAAGG | CTTCATCCTTCTCTGGTTTCT |
| *tpm4a* | AGTGGCCAAACTGGAAA | ATCCAGCTCCTCAGAGAT |
| *acta1b* | CCCAGGTATTGCTGACCGTATG | ACCGATCCATACGGAGTATTTACGC |
| *acta1a* | CAGCTTTGGCTCCAAG | CTGGAAAGTGGACAGAG |
| *actc1a/1c* | TATGAAGTGCGACATTGATATCCGC | ACCAATCCAGACGGAGTACTTGCGT |
| *actc2* | GGCATTATGAAGTGTGACATTGACATTCGT | GCCAATCCAGACAGAGTATTTACGT |
| *actc1b* | GCTACTCTTTCGTGACAAC | ATCTCGTTCTCGAAGTCC |
| *slc8a1a* | CTGAAAGACTCTGTGACTGC | GTCCGCGTACTGATCCT |
| *pln1* | TTCTGCCTCATCCTCATCT | ATGAGAAGCTGGTTACATCAC |
| *ryr2* | GATGTTACAGAACACACTGGA | GAAGCAATCTCCAACAGGAA |
| *serca2* | GCCTGTCATTCTCTTGGAC | CTGCATGGAATACAGAGAGTAG |
| *hsf1* | CCATCGACAGCGGTTTAGAA | GACGCCGCTGAAGAGAAA |
| *bag3* | GCTCAGATCATGGGAGAGA | GATGATTTCTTCAACTGGATGTTC |

### Supplementary Figures

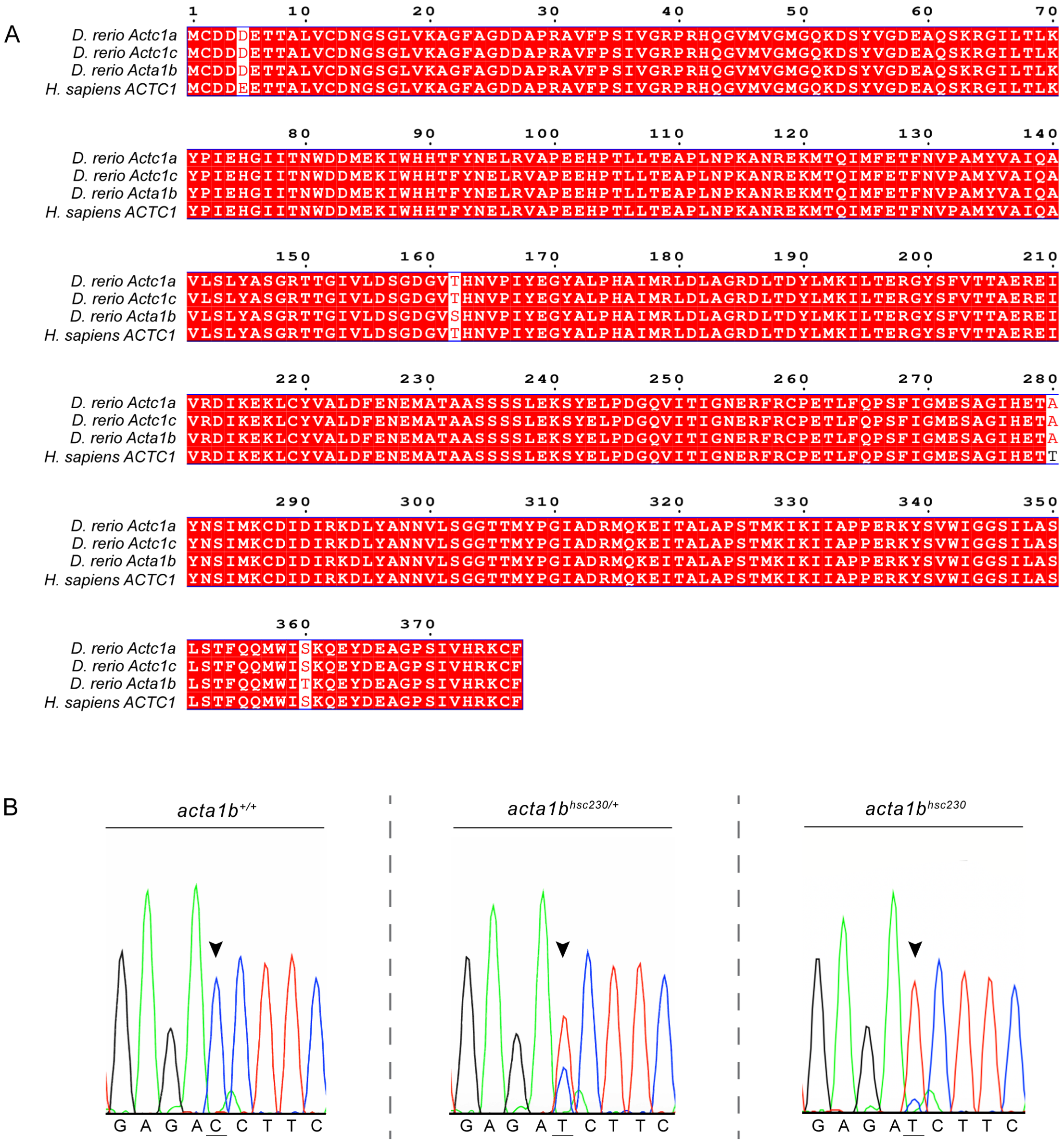

**Supplementary Figure 1. Selection of Acta1b for modeling the human dilated cardiomyopathy-associated T126I mutation in zebrafish.** (A) Protein sequence alignment showing human (ACTC1) and the three primary zebrafish cardiac actin candidates, Actc1a, Actc1c, and Acta1b. (B) Confirmation of *acta1b* c.383C>T point mutation in zebrafish, corresponding to the p.T126I amino acid change.

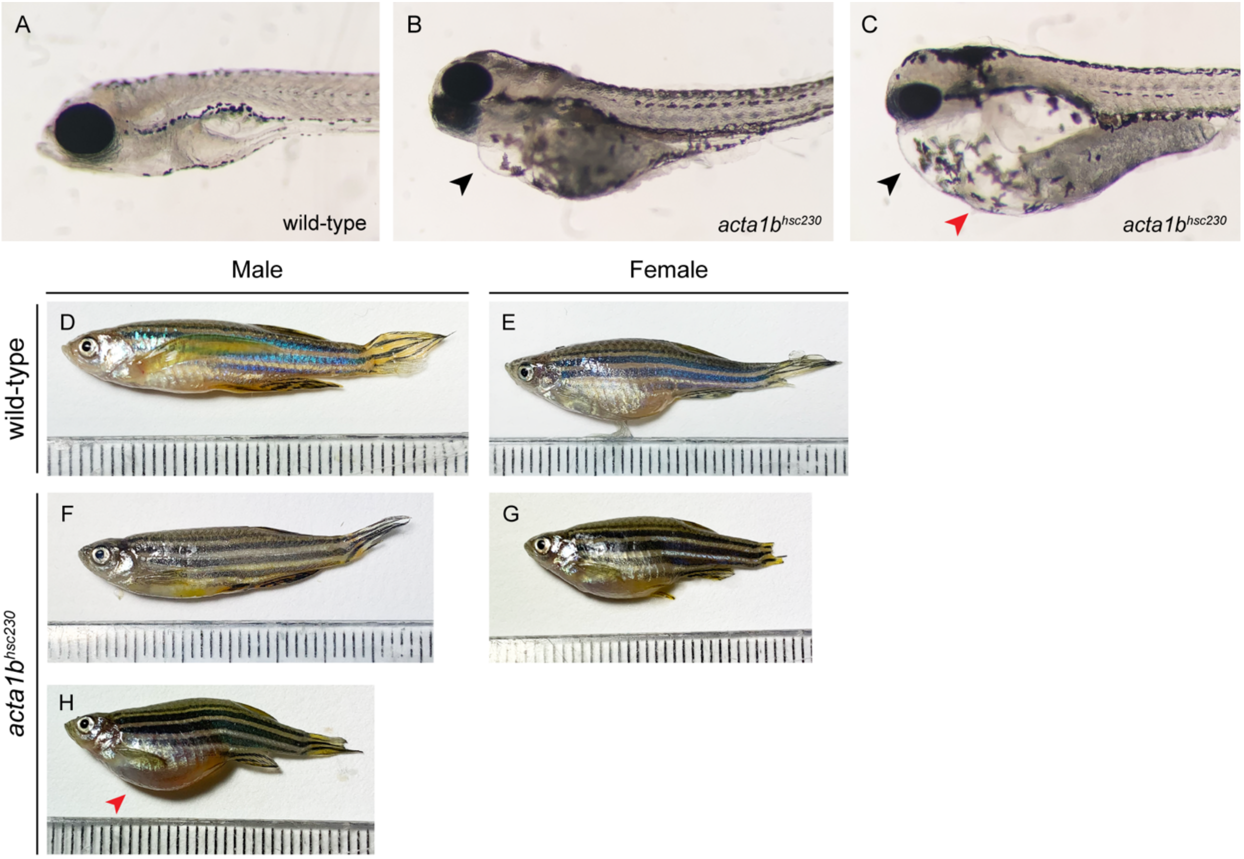
A
wild-type
B
acta1bhsc230
C
acta1bhsc230
Male
Female
wild-type
D
E
acta1bhsc230
F
G
H

**Supplementary Figure 2. Phenotypic comparisons of larval and 6-month-old adult zebrafish.** Lateral views of 7 days-post-fertilization (dpf) (A) wild-type and (B,C) *acta1b*$^{hsc230}$ (B,C) mutants exhibiting varying degrees of phenotypic severity (black arrowhead, pericardial edema; red arrowhead, body edema). Lateral views of 6-month-old adults comparing wild-type males and females (D,E) with *acta1b*$^{hsc230}$ mutant males and females (F-G). The red arrowhead indicates a large body edema present in a male *acta1b*$^{hsc230}$ adult.

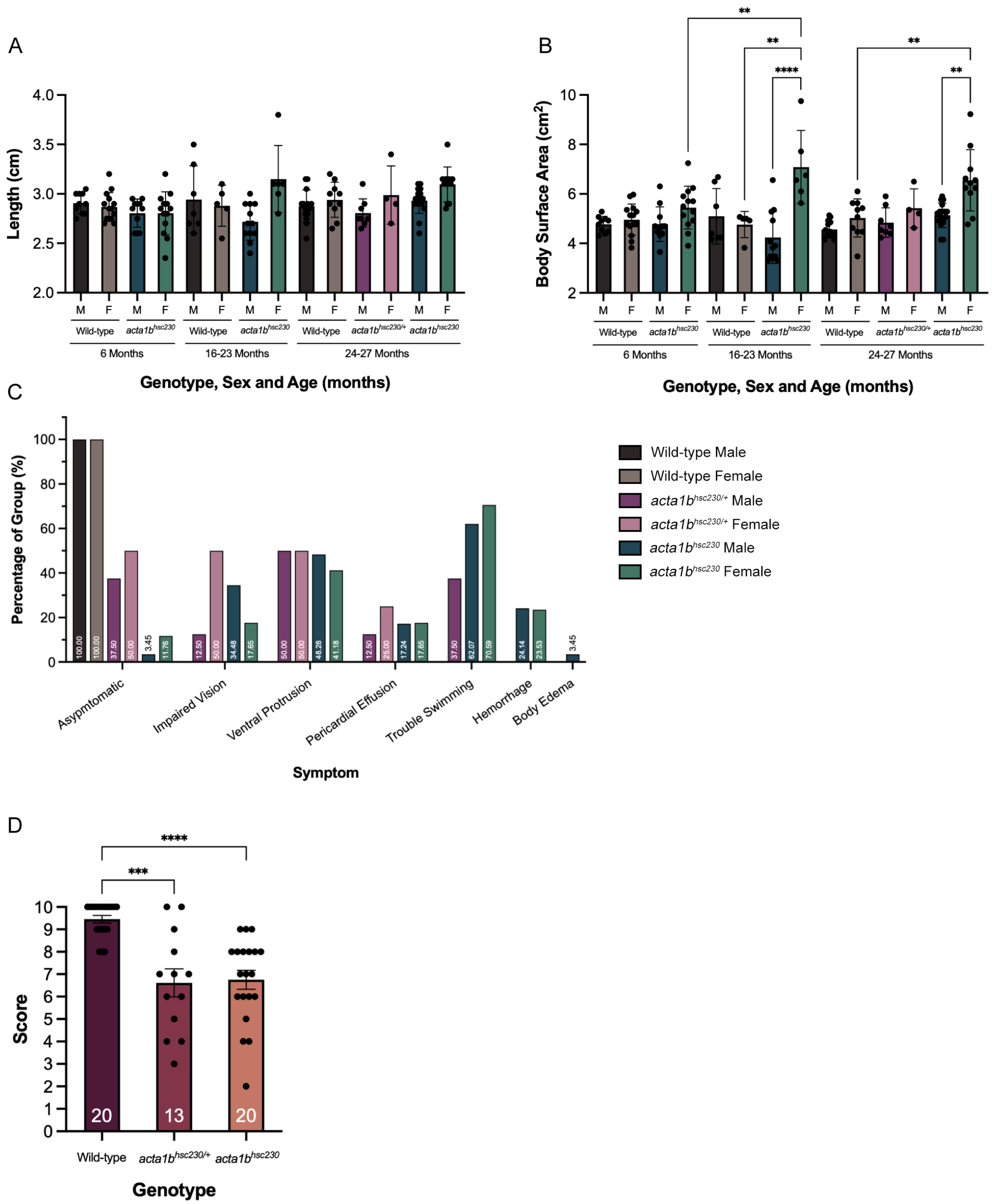
A
Length (cm)
Wild-type
acta1b hsc230
acta1b hsc230/+
6 Months
16-23 Months
24-27 Months
Genotype, Sex and Age (months)
B
Body Surface Area (cm2)
Genotype, Sex and Age (months)
C
Percentage of Group (%)
Asypmtomatic
Impaired Vision
Ventral Protrusion
Pericardial Effusion
Trouble Swimming
Hemorrhage
Body Edema
Symptom
Wild-type Male
Wild-type Female
acta1b hsc230/+ Male
acta1b hsc230/+ Female
acta1b hsc230 Male
acta1b hsc230 Female
D
Score
Wild-type
acta1b hsc230/+
acta1b hsc230
Genotype

**Supplementary Figure 3. Longitudinal analysis of *acta1b*$^{hsc230}$ zebrafish length, body surface area and phenotypes at 16-27 months, with cardiovascular frailty assessment at 24-27 months.** (A) Lengths, (B) body surface area (BSA; 8.46 x [weight]$^{0.66}$), and (C) phenotypes of *acta1b*$^{hsc230}$ mutants from 16-27 months. (D) Frailty scores of wild-type and heterozygous and homozygous *acta1b*$^{hsc230}$ mutants in a test designed to parallel human cardiovascular frailty assessments. Individual values are represented as black dots in panels A,B and D. Total number of individuals tested in each group is indicated by the white number at the bottom of each bar in panel D. Group scores are shown as mean ± SEM. *, $p<0.05$ by two-way ANOVA with Tukey's post hoc multiple comparison test.

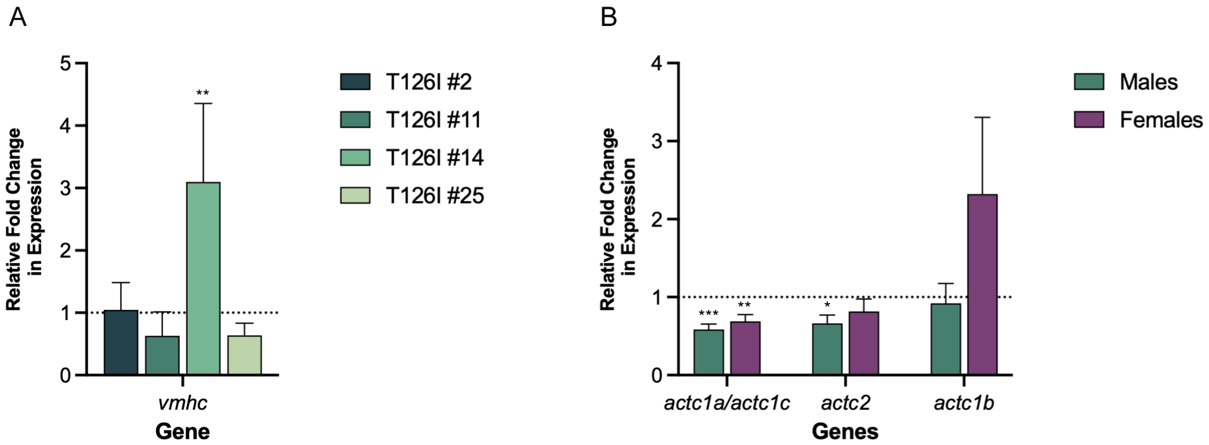
A
Relative Fold Change in Expression
5
4
3
2
1
0
**
vmhc
Gene
T126I #2
T126I #11
T126I #14
T126I #25
B
Relative Fold Change in Expression
4
3
2
1
0
***
**
*
actc1a/actc1c
actc2
actc1b
Genes
Males
Females

**Supplementary Figure 4. Expression of *vmhc* levels in individual mutant males and relative expression of actin paralogues in *acta1b*$^{hsc230}$ mutants at 6 months.** (A) Relative fold change in expression of ventriclular myosin heavy chain (*vmhc*) across four individual *acta1b*$^{hsc230}$ mutant males compared to sex-matched wild-type zebrafish (dotted line indicates wild-type expression level). (B) Relative expression levels of actin paralogues (*actc1a/actc1c, actc2*, and *actc1b*), in 6-month-old male and female *acta1b*$^{hsc230}$ mutants, normalized to their respective sex-matched wild-type controls. Expression values are presented as mean + SEM. *, $p<0.05$ by t-test with unequal variance.
